# Structural basis for gene silencing by siRNAs in humans

**DOI:** 10.1101/2024.12.05.627081

**Authors:** Sucharita Sarkar, Luca F. R. Gebert, Ian J. MacRae

## Abstract

Small interfering RNAs (siRNAs) guide mRNA cleavage by human Argonaute2 (hAgo2), leading to targeted gene silencing. Despite their laboratory and clinical impact, structural insights into human siRNA catalytic activity remain elusive. Here, we show that disrupting siRNA 3’-end binding by hAgo2 accelerates target cleavage and stabilizes its catalytic conformation, enabling detailed structural analysis. A 3.16 Å global resolution cryo-EM reconstruction reveals that distortion of the siRNA–target duplex at position 6 allows target RNA entry into the catalytic cleft and shifts Lysine-709, a previously unrecognized catalytic residue, into the active site. A pyrimidine at target nucleotide t10 positions another unrecognized catalytic residue, Arginine-710, for optimal cleavage. Expansion of the guide-target duplex major groove docks the scissile phosphate for hydrolysis and subsequent groove compression after position 16 permits target RNAs to exit the catalytic cleft. These findings reveal how hAgo2 catalyzes siRNA target hydrolysis, providing a high-resolution model for therapeutic design.

## Introduction

siRNAs are small RNA duplexes that direct potent gene knockdown when introduced into mammalian cells (Elbashir et al., 2001). Over the past two decades, siRNAs have become a critical tool in molecular biology, enabling gene silencing across thousands of studies. siRNAs also hold promise as therapeutic agents for diseases driven by aberrant gene expression, including cancer, viral infections, and genetic disorders. In the last six years, six siRNA-based therapies have entered the clinic, with many more in development. siRNA therapies offer several advantages, including potent and sustained gene knockdown, low side effects, and ease of adaptation to new targets, highlighting their potential as a cornerstone of precision medicine (Jadhav et al., 2024; Tang and Khvorova, 2024).

At the molecular level, siRNAs are loaded into Argonaute 2 (Ago2), where one strand, the guide, is retained while the other is discarded (Leuschner et al., 2006; Martinez et al., 2002). The guide RNA directs hAgo2 to complementary sequences in target mRNAs, which are cleaved by hAgo2’s endonuclease activity (Liu et al., 2004). Efforts to produce potent siRNAs have identified sequence features and chemical modifications that enhance siRNA stability and knockdown efficiency (Foster et al., 2018; Parmar et al., 2016; Reynolds et al., 2004; Vert et al., 2006), as well as guide-target pairing parameters that influence target mRNA cleavage (Becker et al., 2019; Wang and Bartel, 2024). However, most of this work has been empirical, with limited structural insights to guide optimization.

Recently, moderate-resolution cryo-EM structures of *Arabidopsis* Ago10 (AtAgo10) (Xiao et al., 2023) and hAgo2 (Mohamed et al., 2024), bound to guide and fully complementary target RNAs, have shed light on the domain movements required for catalysis. While these studies provided valuable insights into how eukaryotic Argonautes adapt their shape to engage target RNAs, the resolutions achieved were inadequate to visualize and accurately model individual RNA bases and numerous protein side chains. Thus, despite the growing clinical relevance of siRNAs, a mechanistic basis for optimizing their activity remains elusive.

### siRNA 3’-End Release Is a Limiting Step in Target Slicing

To visualize the gene-silencing mechanism employed by siRNAs in humans, we examined samples of the hAgo2-siRNA complex engaged with a complementary target RNA using cryo-EM and single-particle analysis. Initial efforts resulted in heterogeneous particles and low-resolution reconstructions lacking clear RNA density. This outcome aligns with single-molecule studies reporting hidden hAgo2 conformations involving dynamic release and re-anchoring of the siRNA 3’-end during target engagement (Willkomm et al., 2022). To favor the catalytic conformation, we engineered hAgo2 mutant proteins designed to facilitate 3’-end release by removing interactions involved in guide 3’-end binding (Fig. 1a). Two different 3’ end-binding mutants were produced (F294A and R315V/H317A). Both displayed an initial burst of target cleavage activity about 7 times faster than wild-type hAgo2, followed by similar steady-state cleavage rates (Fig. 1b and S1). This finding indicates that 3’-end release is rate-limiting in the first round of catalysis (Fig. 1c). Moreover, when catalysis was inhibited by excluding divalent cations from the reaction mixture, both mutants yielded cryo-EM reconstructions featuring a fully formed siRNA-target duplex (Fig. S2), enabling us to determine the structure of the catalytic conformation at a global resolution of 3.16 Å (Fig. 2, S3-S6, and Table S1).

**Fig. 1.**
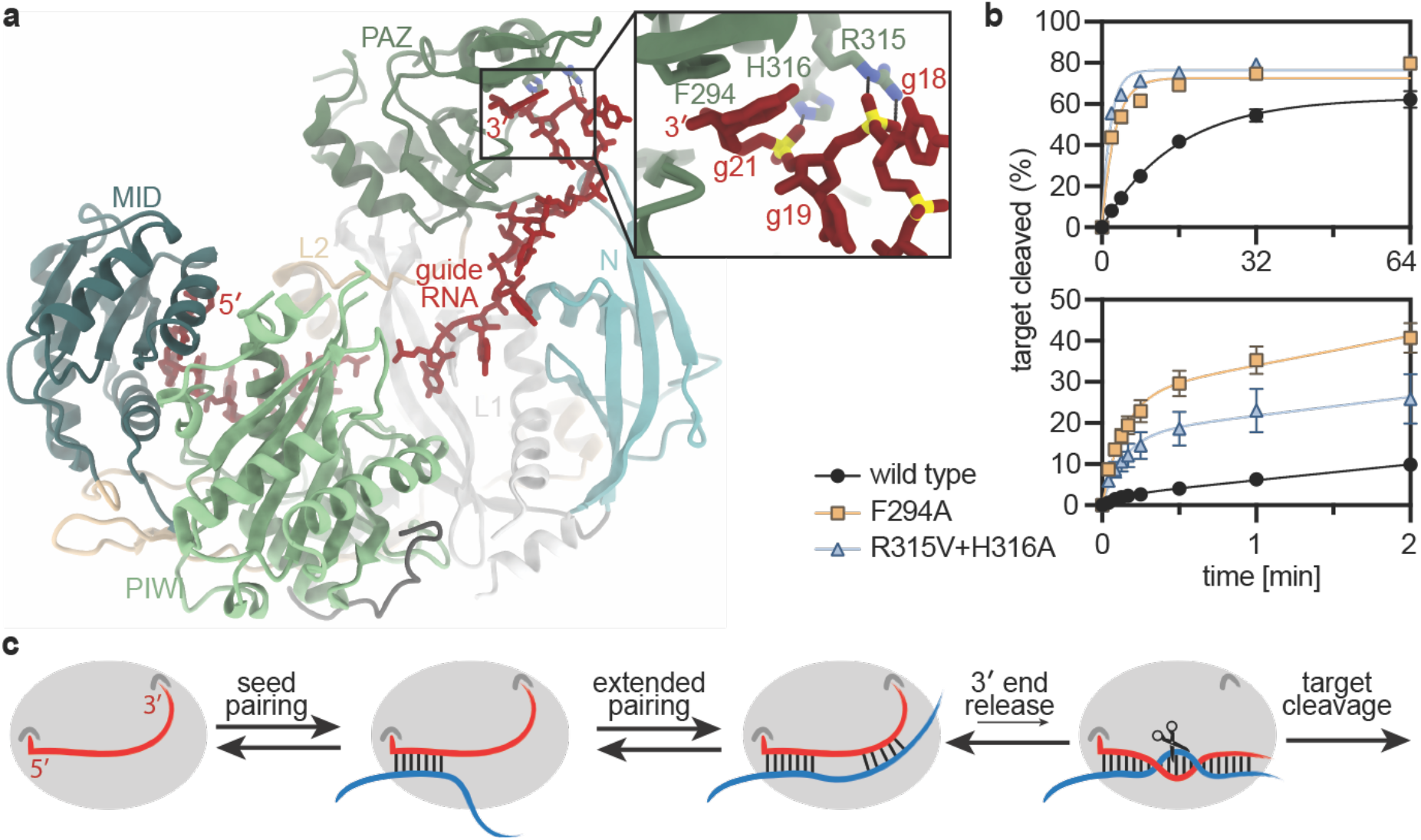
Guide 3’-end release favors the catalytic conformation. **a**. Structure of the hAgo2-guide RNA complex (PDB 4W5N) with major domains labeled. Inset shows residues mutated to weaken interactions with the guide 3’-end. **b**. Cleavage of a target RNA over time by wild type and 3’-end binding hAgo2 mutants under single turnover (top panel) and multiple turnover (bottom panel) conditions. Data points are the mean values of 3 independent experiments; error bars indicate SEM. **c**. Proposed kinetic scheme wherein 3’-end release is rate-limiting toward achieving the catalytic conformation and target cleavage.

**Fig. 2.**
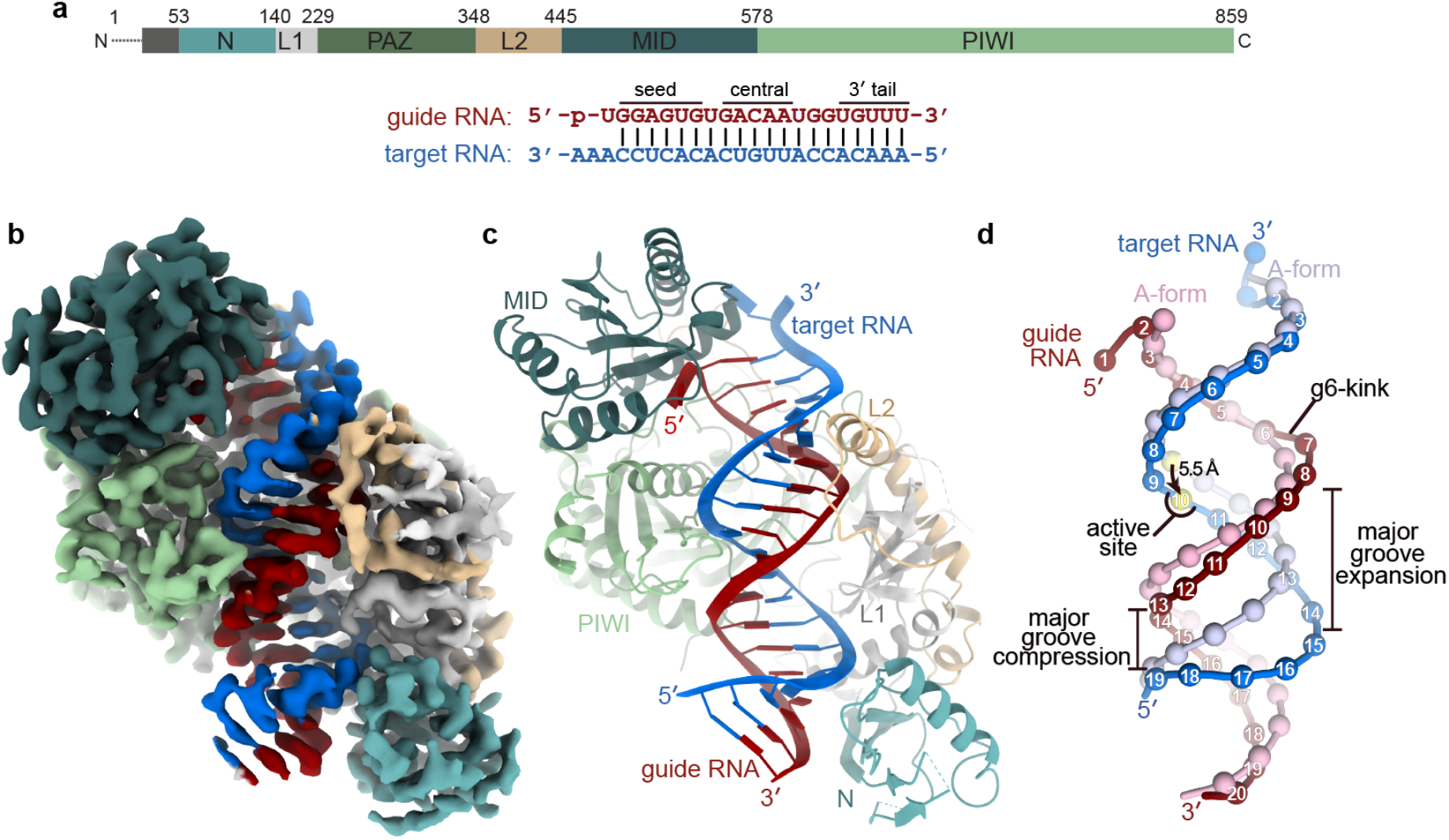
The catalytic conformation of human Ago2. **a**. Schematics of hAgo2, guide RNA, and target RNA primary sequences. **b**. Reconstruction of the R316V+H317A hAgo2 mutant engaged with guide and target RNAs (without supplemented divalent cations necessary for catalysis). **c**. Cartoon representation of the hAgo2-guide-target atomic model. **d**. Superposition of the hAgo2-bound guide-target duplex (dark red and blue) onto an ideal A-form duplex (pink and light blue). RNAs were aligned to the seed region, which is anchored onto hAgo2(Sheu-Gruttadauria et al., 2019a). Spheres denote phosphate positions, numbered from the guide 5’ end; target phosphates match the numbering of their paired guide nucleotides.

### Structure of the hAgo2 Catalytic Conformation

The reconstruction of hAgo2 in its catalytic conformation reveals density for all protein domains except the PAZ domain, which contains the guide 3′ end-binding site and is disordered (Fig. 2a–c). This observation suggests that release of the guide RNA 3′ end uncouples PAZ domain movements from the rest of the protein. Notably, a well-defined guide-target RNA duplex is visible traversing the central cleft of hAgo2 (Fig. 2b). In their recent lower-resolution hAgo2-guide-target structure, Mohamed et al. observed moderate deviations from A-form geometry in the duplex but did not address potential functional implications of these distortions (Mohamed et al., 2024). In our higher-resolution reconstruction, these helical distortions are clearly defined and include a kink near the end of the seed region (g6-kink), an expanded major groove in the guide central region, and a compressed major groove in the 3′ tail region (Fig. 2d). Together, these structural features facilitate the passage of the guide-target duplex through the hAgo2 cleft, position the scissile phosphate, and align catalytic residues for efficient target RNA cleavage.

### The g6-kink Couples Guide-Target Pairing to Target Cleavage

The g6-kink is a bend in the guide RNA backbone between positions g6 and g7 (Fig. 2d). As previously noted (Wang and Bartel, 2024), the g6-kink involves guide nucleotide g6 adopting a C2’-endo sugar pucker conformation (Fig. S7a). The kink is stabilized by hydrophobic interactions between Ile-365 and the sugar face of the guide-target duplex around positions 6 and 7 (Fig. S7b). Ile-365 resides on helix-7, a mobile structural element that shifts as the hAgo2 central cleft widens to accommodate guide-target pairing (Fig. 3a) (Klum et al., 2017; Schirle et al., 2014; Sheu-Gruttadauria et al., 2019b). Consequently, the severity of the g6-kink correlates with the extent of base pairing between the guide and target RNAs, providing an internal sensor of pairing across the guide-target duplex (Fig. 3b-c).

**Fig. 3.**
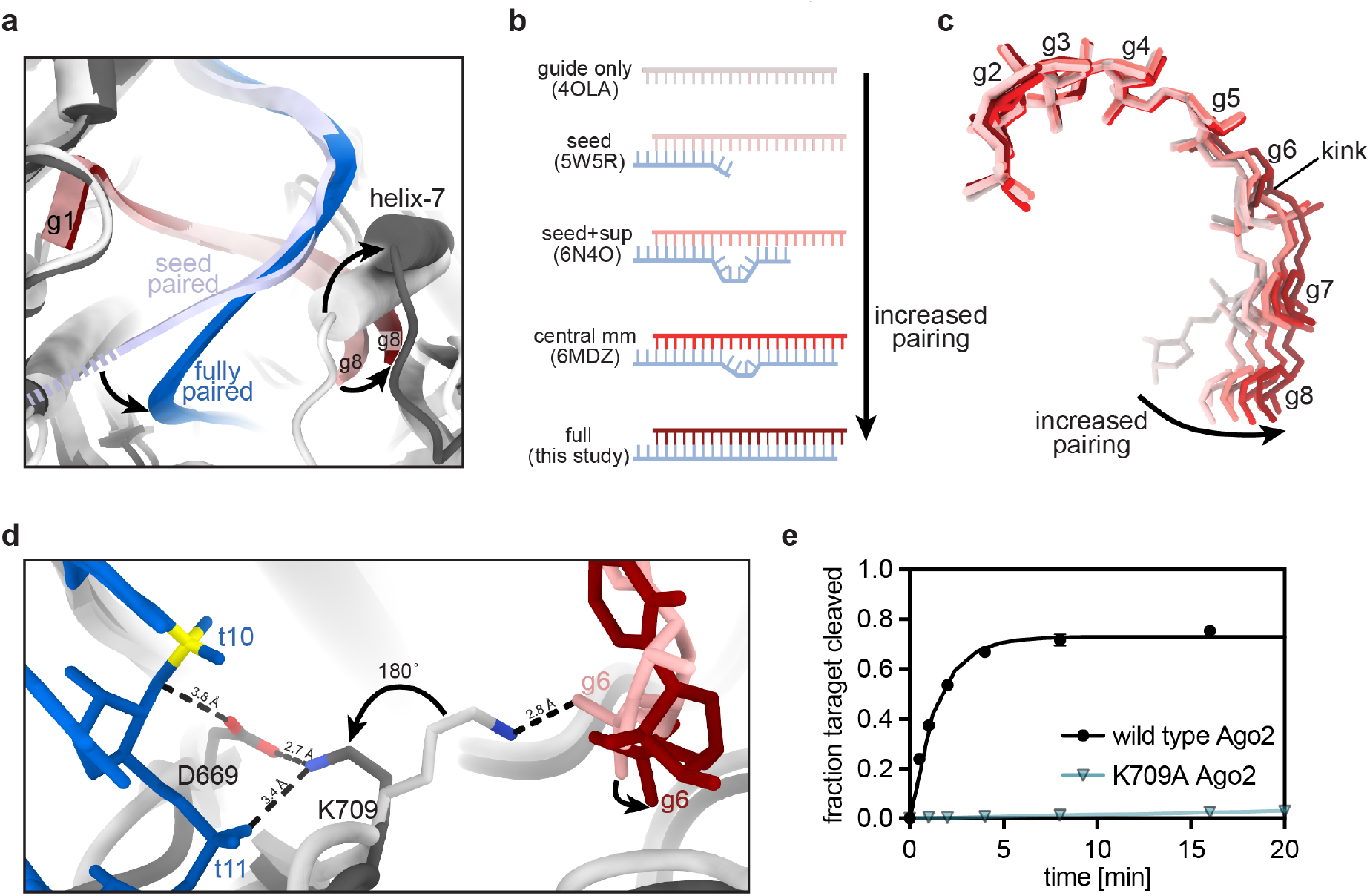
The g6-kink senses target pairing and licenses cleavage. **a**. Superposition of seed-paired hAgo2 (light colors, 5W5R) onto the fully paired hAgo2 structure (dark colors). Arrows indicate conformational changes moving from seed to fully paired. **b**. Base pairing schematics for published hAgo2 structures. **c**. Superposition of sugar-phosphate backbone of seed region nucleotides from hAgo2 structures listed in panel b. Guide RNAs are colored as in b. **d**. Superposition of seed-paired (light colors) and fully paired (dark colors) structures. The extended g6-kink correlates with the repositioning of K709 into the active site. Arrows indicate the directions of change from seed-paired to fully paired. P atom in scissile phosphate (t10) colored yellow. **e**. Fraction of target RNA cleaved as a function of time by wild type hAgo2 and K709A hAgo2. Data points are the mean values of 3 independent experiments; error bars indicate SEM.

The g6-kink enables target cleavage in two ways. First, the presence of the kink alters the trajectory of the guide-target duplex after the seed region. This adjustment avoids steric clashes with the PIWI domain as the guide-target duplex enters the hAgo2 central cleft (Fig. 3a) and helps position the target strand in the active site (Fig. 2d). Second, the kink moves the g6 phosphate, which forms a salt linkage to the amine group lysine-709 (K709) in guide-only and seed-paired hAgo2 structures (Fig. 3d). Upon extended pairing, the K709 side flips into the active site, where it contacts the t11 phosphate and the catalytic residue Aspartate-669 (D669). Previous hAgo2 structures did not reveal the K709-D669 interaction because seed-pairing alone does not induce a kink large enough to disrupt the g6–K709 salt link (Schirle et al., 2014), and the structures with pairing beyond the seed used a D669A mutant (Mohamed et al., 2024; Sheu-Gruttadauria et al., 2019a; Sheu-Gruttadauria et al., 2019b; Xiao et al., 2023).

Inspection of eukaryotic, bacterial, and archaeal Argonaute structures revealed the Lys-Asp pair analogous to K709-D669 is a deeply conserved feature of the Argonaute active site (Fig. S8). Molecular mechanics simulations indicate that the equivalent Asp residue in RNase H donates a proton to the 2′ oxygen leaving group on the 5’ cleavage product (Rosta et al., 2011), suggesting K709 may contribute to the final chemical step of target RNA hydrolysis. Supporting this model, the K709A hAgo2 mutant retains target-binding activity but cleaves target RNAs ∼500x slower than wild-type hAgo2 (Fig. 3e).

The combined results suggest that the g6-kink acts as a sensor for guide-target pairing that licenses target cleavage. Upon extended guide-target pairing, the g6-kink shifts the position of the g6 phosphate enough to liberate K709. K709 can then flip into the active site, where it positions the target strand through interactions with t11 and enables D669 to catalyze hydrolysis (Fig. 3d). Features that increase guide RNA flexibility in the proximity of the g6-kink such as AU pairing at position 7 (Reynolds et al., 2004; Wang and Bartel, 2024) and glycol nucleic acid (GNA) substitutions (Egli et al., 2023; Schlegel et al., 2017) in Enhanced Stabilization Chemistry plus (ESC+) siRNAs (Schlegel et al., 2022) have been shown to enhance hAgo2 activity, and a g6G:t6G mismatch makes the otherwise inactive zebrafish Ago2 active (Chen et al., 2017).

### Scissile Phosphate Positioning Requires Major Groove Expansion from t10–t14

Modeling an ideal A-form RNA duplex onto the 5’ end of the seed region, which is fixed on hAgo2 (Sheu-Gruttadauria et al., 2019a), reveals a ∼5.5 Å shift in the target RNA scissile phosphate relative to A-form (Fig. 2d). This shift is caused by the g6-kink and expansion of the major groove near the center of the guide-target-duplex (Fig. 4a). Major groove expansion appears to be driven by interactions between hAgo2 and target nucleotides t10-t14, which dock in an extended conformation into a narrow channel on the surface of the hAgo2 PIWI domain (Fig. 4b). Notably, target cleavage rates are particularly sensitive to central region mismatches, which can either accelerate or inhibit catalysis (Becker et al., 2019). We propose that docking t9–t14 promotes a major groove expansion that pulls the scissile phosphate (t10) into the active site and exaggerates local distortions associated with base-pair mismatches, increasing hAgo2’s sensitivity to mispairing the central region. Moderate stability in the guide central region is a general feature of effective siRNA sequences (Khvorova et al., 2003), and locked nucleic acid (LNA) substitutions, which increase duplex stability and rigidity, are poorly tolerated at g10, g12, and g14 (Elmen et al., 2005).

**Fig. 4.**
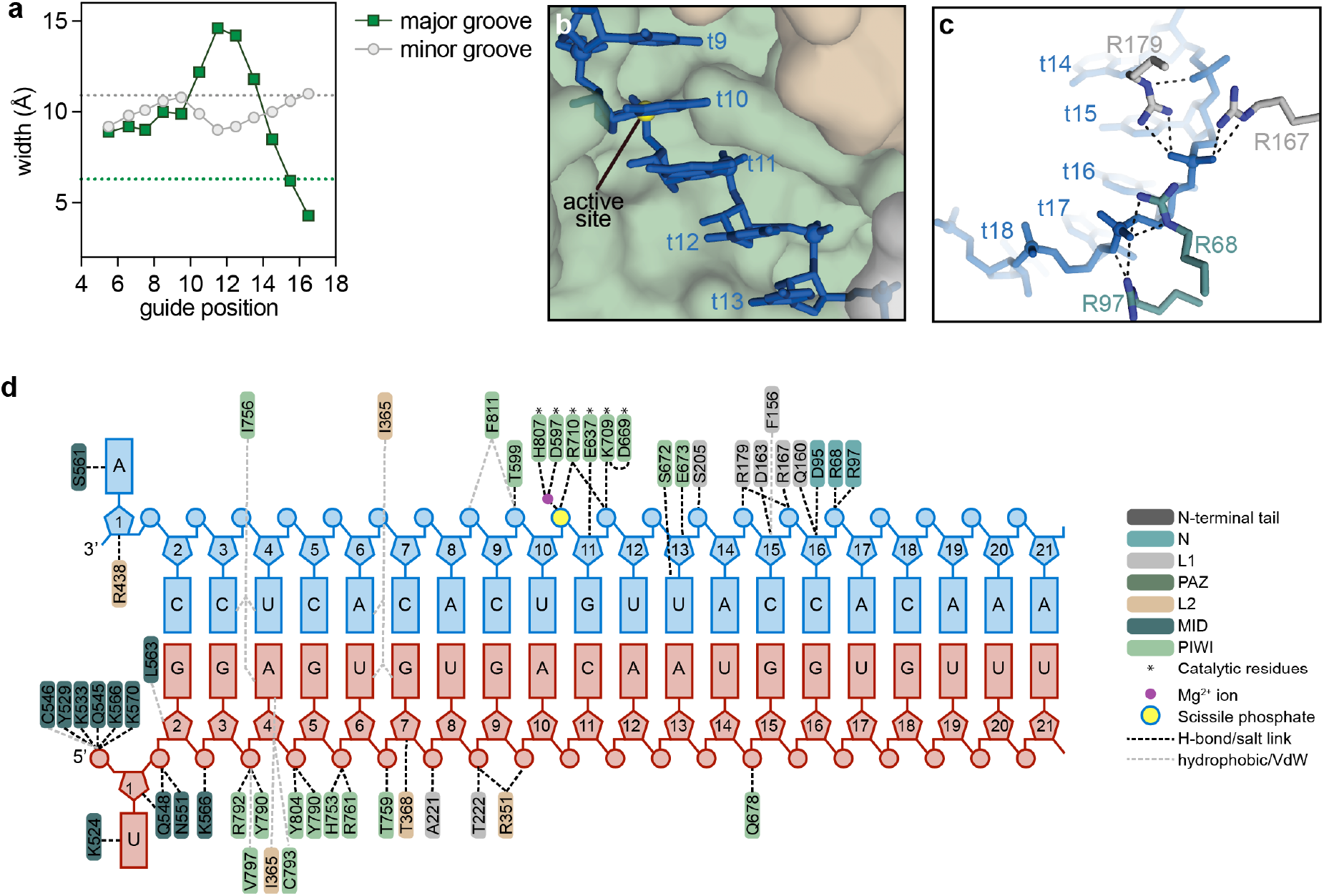
Duplex docking requires modulation of guide-target major groove. **a**. Refined major groove (green) and minor groove (grey) widths along the guide-target duplex. Dashed lines indicate ideal groove widths. **b**. Target nucleotides t9-t13 dock in an extended conformation along the surface of hAgo2. **c**. Salt linkages between the t14-t16 phosphate backbone and arginine residues from the N (green) and L2 (gray) domains dock the end of the guide-target duplex in the central cleft. **d**. Schematic of the specific contacts between hAgo2 and the guide (red) and target (blue) RNAs observed in the experimental map.

### Docking the Guide-Target Duplex Involves Major Groove Compression after t16

The backbone phosphates of t14-t16 are extensively contacted by residues from the hAgo2 N and L1 domains, stabilizing the docked guide-target duplex within the central cleft (Fig. 4c-d). These contacts illuminate how the N-domain contributes to target cleavage and mark the end of hAgo2’s engagement with the guide-target duplex—after position 16, no specific contacts between the protein and RNA are observed. However, the protein nonetheless exerts influence on the RNA as the major groove narrows sharply at position 16 to avoid clashes with the N domain and allow the duplex to exit the central cleft (Fig. 2c-d, 4a, S7c-d). This explains why guide-target mismatches beyond position 16 are often well-tolerated, or even enabling, for target cleavage (Becker et al., 2019). Additionally, the compression of the major groove explains why AU base pairs, which promote major groove narrowing (Marin-Gonzalez et al., 2020), at position 17 enhance cleavage rates (Wang and Bartel, 2024).

### Arg-710 Accelerates Target Cleavage and is Sensitive to t10 Identity

Early studies examining potent siRNA sequences found that an A nucleotide at g10 (g10A) correlates with elevated knock-down efficiency (Reynolds et al., 2004; Vert et al., 2006). Similarly, kinetic studies show that guide sequences with a g10 purine direct faster target cleavage than equivalent guides with a g10 pyrimidine (Wang and Bartel, 2024). Examination of the hAgo2 structure shows that no part of the protein is positioned to interrogate the identity of g10. However, the guanidinium group of Arginine-710 (R710) resides near the nucleobase of t10 (the target nucleotide paired with g10). R710 also directly interacts with the scissile phosphate, suggesting a role in catalysis (Fig. 5a). Indeed, mutating R710 to alanine reduced target cleavage by about 20-fold (Fig. 5b). Slower cleavage by an R710A mutant hAgo2 using a different guide RNA sequence was also reported by Mohamed and coworkers (Mohamed et al., 2024).

**Fig. 5.**
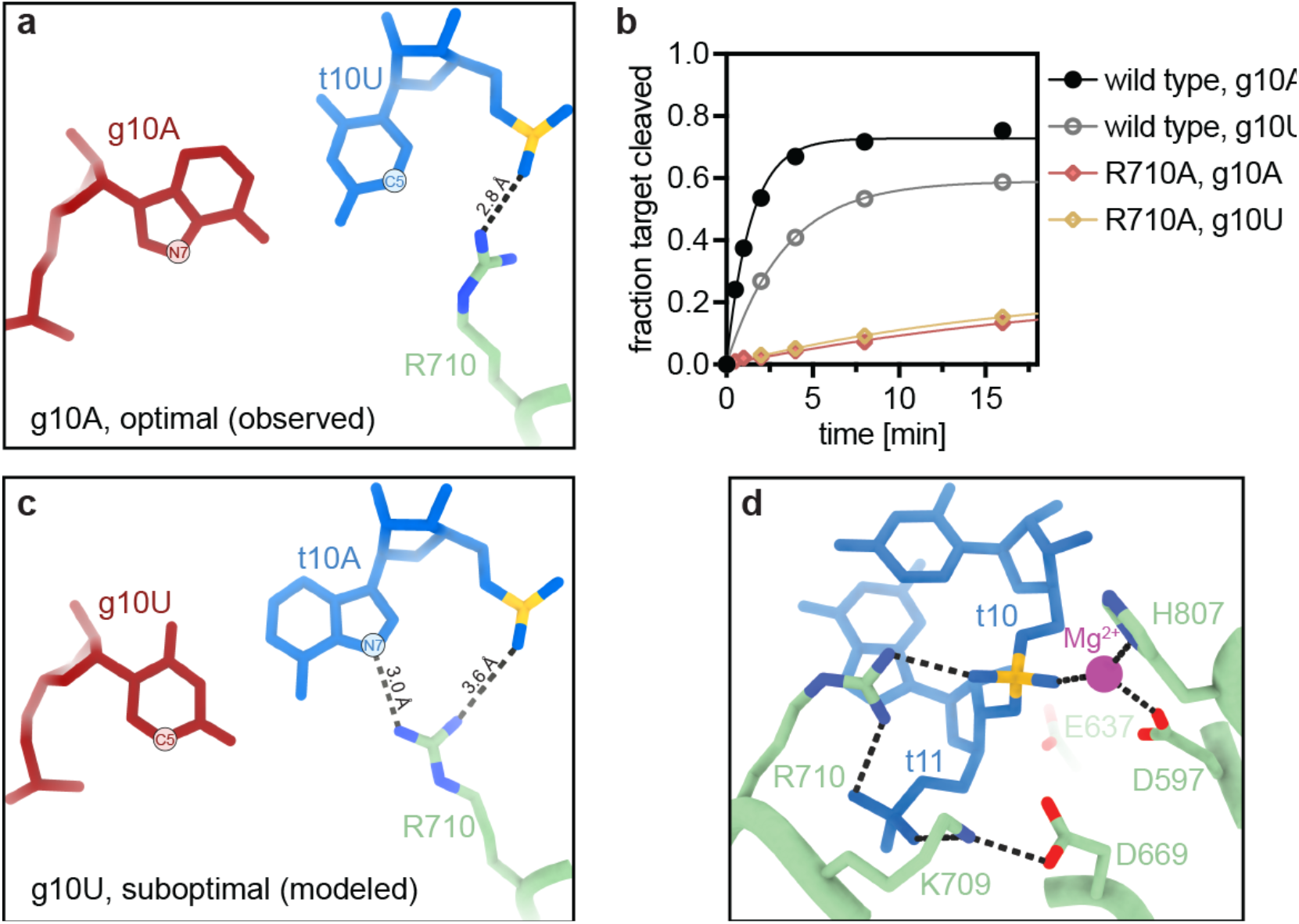
Active site optimization. **a**. Observed structure with R710 forming a salt linkage with a non-bridging oxygen on the scissile phosphate. **b**. Single turnover cleavage kinetics of wild-type hAgo2 and the R710A hAgo2 mutant using either g10A (optimal) or g10U (suboptimal) guide RNAs. Data points are the mean values of 3 independent experiments; error bars indicate SEM. **c**. Modeled structure of the suboptimal g10U conformation in which R710 interacts with the N7 atom of t10 purines. **d**. The updated model of the hAgo2 active site includes the previously established catalytic tetrad (D597, E637, D669, and H807) (Liu et al., 2004; Nakanishi et al., 2012) plus residues identified in this study: K709, proposed to facilitate protonation of the leaving group, and R710, proposed to stabilize charge on the transition states. A single co-purifying Mg^2+^ ion was also observed in our density. A second, more labile, Mg^2+^ is expected to bind between D597, D669, and E637(Sheng et al., 2014).

In our structure, t10 is a U nucleotide, making g10 an A, the pairing predicted to be optimal for catalysis (Reynolds et al., 2004; Vert et al., 2006; Wang and Bartel, 2024). Consistent with this expectation, switching the nucleotide identities at position 10 (to g10U and t10A) reduced the cleavage rate by approximately 2.5-fold (Fig. 5b). The C-H edge of the t10U is positioned too far from R710 to indicate a clear positive interaction (3.7 Å between the C5 atom of t10U and the Nη1 atom of R710). By contrast, modeling an A or G nucleotide at t10 presents the N6 amine of the purine Hoogstein face for hydrogen bonding to R710 (Fig 5c), a shift that would move Arg-710 away from the scissile phosphate.

Based on these observations, we propose that R710 promotes target cleavage by stabilizing charged intermediates and transition states along the hydrolysis reaction pathway. In this model, R710 accelerates catalysis but is not strictly required for activity. A similar role was proposed for Arg-116 in *E. coli* alkaline phosphatase, which uses a two-metal mechanism akin to hAgo2 (Kim and Wyckoff, 1991). R710 may also help position the target RNA through interactions with the t11 phosphate. We further propose that the activity of R710 is influenced by the identity of the t10 nucleotide, with purines at t10 promoting an alternative, less active conformation (Fig. 5c). Supporting this hypothesis, the R710A hAgo2 mutant does not display the same t10 nucleotide preference as wild-type hAgo2 (Fig. 5b).

## Discussion

The data presented here offer the first detailed view of the catalytic conformation of human Ago2, providing mechanistic insights into how siRNAs drive the silencing of target mRNAs. Notably, our structure differs from the model recently reported by Mohamed and coworkers (Mohamed et al., 2024). While they propose that an exaggerated positional shift in the N domain is critical for allowing the RNA full access to the central channel, along with several RNA-protein contacts that we do not observe, our structure shows that the RNA can instead be accommodated by compaction of the major groove in the 3’ tail region (Fig. S9). Thus, an exaggerated N domain shift is not a formal requirement for full RNA access to the central cleft. Alternatively, we suggest that differences between the models indicate that target RNA catalysis may involve multiple related conformations, the distribution of which is dictated by features within the guide-target duplex. Considering that the guide sequence used by Mohamed and coworkers (miR-7) drives faster cleavage rates than the guide used in our structure (miR-122) (Wang and Bartel, 2024), a direct comparison of the maps and models from both studies will be informative.

The findings presented here provide mechanistic explanations for many empirically derived siRNA optimizations, including sequence preferences at positions 7, 10, and 17 (Reynolds et al., 2004; Vert et al., 2006; Wang and Bartel, 2024), the stimulatory effects of g6 mismatches and GNA substitutions (Chen et al., 2017; Egli et al., 2023; Schlegel et al., 2017), the negative impact of LNAs in the guide central region (Elmen et al., 2005), and how 3’ tail mismatches can stimulate catalysis (Becker et al., 2019). More importantly, the data elucidate the specific contacts that dock target RNAs for cleavage and reveal how RNA-protein interactions modulate the conformation and activity of previously uncharacterized catalytic residues. These results provide a larger and more dynamic model of the hAgo2 active site (Fig. 5d) and a rational foundation for the next generation of siRNA design.

## Data Availability

The cryo-EM map for the hAgo2 R315V/H316A-guide-target RNA complex was deposited in the EMDB under accession ID EMD-46888 and the corresponding atomic model coordinates were deposited in the Protein Data Bank under accession code PDB ID 9DHX.

## Limitations of this study

The hAgo2 protein used for structure determination here contains two mutations in the guide 3’-end binding pocket (R315V/H316A) and four mutations at positions previously shown to be phosphorylated (Schirle and MacRae, 2012). All of these mutated residues were disordered in our reconstruction: density for the entire PAZ domain, which includes the 3′-end binding pocket, was almost entirely lacking, and, as in previous hAgo2 structures, the mobile element containing the phosphorylation sites was not observed. Although these regions were unresolved, we cannot exclude the possibility that the mutations affect protein or RNA structure in unforeseen ways. Additionally, 3D reconstruction by single-particle analysis involves excluding a significant fraction of observed particles, potentially omitting distinct alternate conformations associated with target RNA catalysis. Finally, our structural analysis was conducted without exogenous Mg^2^+to prevent target cleavage. Because the active site likely contains a second, more liable Mg^2+^ binding site (Sheng et al., 2014) that is vacant in our structure, the conformations of catalytic side chains may shift during the catalytic processes.

## Acknowledgments

This research was funded by NIGMS grant R35GM127090 and through a sponsored research agreement with Eli Lilly and Company.

## Methods

### Preparation of recombinant hAgo2-guide RNA complexes

Human Ago2 proteins were expressed in Sf9 cells using the baculovirus system according to a previously published protocol (Sheu-Gruttadauria and MacRae, 2018). Pelleted cells, resuspended in lysis buffer (50 mM NaH2PO4 pH 8, 100 mM NaCl, and 0.5 mM TCEP), were lysed by passing them three times through an M-110P microfluidizer (Microfluidics). The lysate was clarified by centrifugation at 12,000 g for 15 min, followed by an initial purification by Ni affinity (Ni NTA agarose, Qiagen) combined with Micrococcal nuclease (Takara) treatment in imidazole wash buffer (50 mM Tris-HCl pH 8, 300 mM NaCl, 20 mM imidazole, 5 mM CaCl2, and 0.5 mM TCEP). hAgo2 was loaded with a synthetic guide RNA oligonucleotide bearing a 5’-terminal phosphate (IDT). The loaded hAgo2 RNPs were captured using a biotinylated antisense oligonucleotide (2’-O-Me RNA, IDT) immobilized on high capacity neutravidin agarose (Thermo Fisher Scientific), washed with 3 different buffers (first: 30 mM Tris-HCl pH 8, 0.1 M KOAc, 2 mM Mg(OAc)2, 0.02% CHAPS, 0.5 mM TCEP - second: 30 mM Tris-HCl pH 8, 2 M KOAc, 2 mM Mg(OAc)2, 0.02% CHAPS, 0.5 mM TCEP - third: 30 mM Tris-HCl pH 8, 1 M KOAc, 2 mM Mg(OAc)2, 0.02% CHAPS, 0.5 mM TCEP) and eluted with a biotinylated DNA competitor (IDT). Excess competitor was removed by incubation with high capacity neutravidin agarose (Thermo Fisher Scientific). Finally, residual oligonucleotide contaminants were removed by incubation of the protein with Q Sepharose Fast Flow resin (Cytiva). The protein was buffer exchanged into storage buffer (50 mM Tris-HCl pH 8, 100 mM NaCl, 0.5 mM TCEP), concentrated, aliquoted, flash-frozen in liquid nitrogen, and stored at -80 °C. The concentration was determined on a Nanodrop OneC (Thermo Fisher Scientific) using an extinction coefficient (198370 M^−1^ cm^−1^) calculated for absorbance by hAgo2 and guide RNA at 280 nm.

### Burst and steady-state kinetics

Multiple turnover condition reactions were set up in 30 mM Tris-HCl pH 8.0, 0.1 M KOAc, 2 mM Mg(OAc)2, 0.5 mM TCEP, with a total of 100 nM RNA (of which 1-2 nM was ^32^P-radiolabeled on the target 5’ end), and 0.1 mg/ml Baker’s Yeast tRNA. A zero time point was taken by transferring 10 μl of the mixture into an equal volume of 2x formamide loading buffer (0.05% SDS, 0.5 mM EDTA, 0.025% bromophenol blue, 0.025% xylene cyanol in formamide). Reactions were then started by adding the hAgo2:guide complex to a final concentration of 50 nM. Subsequent time points were taken by transferring 10 μl of the reaction solution into 10 μl 2x formamide loading buffer. The time points were analyzed by 15% acrylamide 8 M urea denaturing polyacrylamide gels in 0.5x TBE. The gels were used to expose a phosphor screen (Cytiva) for visualization. Screens were imaged on a Typhoon phosphorimager (Cytiva) and the signal was quantified using ImageQuant (Cytiva). The data were fit to the burst and steady-state equation as described(Arif et al., 2022) using Prism (GraphPad Software):

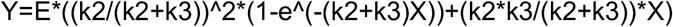

where Y is the amount of product in nM, E is the concentration of active fraction of enzyme, k2 is the slicing rate, and k3 is the product release rate. E was derived from the single turnover kinetics and the nominal enzyme concentration.

### Single turnover kinetics

Single turnover reactions were set up in 30 mM Tris-HCl pH 8.0, 0.1 M KOAc, 2 mM Mg(OAc)2, 0.5 mM TCEP, with a total of 0.2 nM ^32^P-radiolabeled RNA, and 0.1 mg/ml Baker’s Yeast tRNA. A zero time point was taken (into 1 volume formamide loading buffer), and the reactions were started by the addition of the hAgo2:guide complex to a final concentration of 5, 10, or 20 nM. Subsequent time points were taken by transferring 10 μl reaction to 10 μl 2x formamide loading buffer. The time points were analyzed by 15% acrylamide 8 M urea denaturing polyacrylamide gels in 0.5x TBE. The gels were used to expose a phosphor screen (Cytiva) for visualization. Screens were imaged on a Typhoon phosphorimager (Cytiva) and the signal was quantified using ImageQuant (Cytiva). The data were fit to a simple one-phase decay curve in Prism (GraphPad Software). In these experiments, multiple hAgo2 concentrations were used for each condition/mutant to ensure that target cleavage, not target binding, was the observed rate-limiting step. Within each experiment, plots using the different Ago2 concentrations were always indistinguishable from each other. Therefore, all data were combined to create the plots displayed in the manuscript.

### Grid preparation for cryo-EM

The hAgo2 R315V/H316A-guide-target RNA complex was formed at room temperature for 10 minutes in storage buffer (50 mM TRIS, pH 8.0, 100 mM NaCl, and 0.5 mM TCEP) by mixing hAgo2-guide and target RNA at equimolar ratios. Two samples were prepared, yielding final complex concentrations of 0.7 mg/mL and 1.3 mg/mL, respectively.

For grid preparation, 300 mesh R1.2/1.3 UltraAuFoil Holey Gold grids (Quantifoil) were subjected to glow discharge under vacuum conditions for 30 seconds at a current of 15 mA using a Pelco easiGlow 91000 Glow Discharge Cleaning System (Ted Pella Inc.). Subsequently, 4 μL of the sample was applied to each grid surface, followed by blotting with Whatman #1 filter paper (55 mm diameter) for 4 seconds with a blot force of 4 on a Vitrobot Mark IV (Thermo Fisher Scientific) operating at 100% humidity and 4°C. The grids were then plunge-frozen in liquid ethane.

The hAgo2 F294A-guide-target RNA, hAgo2 D669A-guide-target RNA, and hAgo2 WT-guide-target RNA complexes were formed at room temperature for 10 minutes in storage buffer (50 mM TRIS, pH 8.0, 100 mM NaCl, and 0.5 mM TCEP) at a 1:1.5 molar ratio of protein-guide to target RNA, yielding final concentrations of protein-guide at 0.7-1.2 mg/mL and target RNA at 1.1-1.8 mg/mL. The grids were prepared as described above.

### Cryo-EM data acquisition

The cryo-EM data of the hAgo2 R315V/H316A-guide-target RNA complex were collected on a TALOS Arctica transmission electron microscope (Thermo Fisher Scientific) with a field emission gun operating at an accelerating voltage of 200 kV and equipped with a Falcon 4i Direct Electron Detector. The cryo-EM datasets were acquired using the EPU automated data collection software (Thermo Fisher Scientific). The movies were collected over an 3.2 s exposure resulting in a total cumulative electron exposure of 40 e^-^Å^-2^. The datasets were collected with C2 aperture 50 μm, magnification of 190,000x, corresponding to 0.74 Å per pixel on the detector with defocus range of -1.0 to -3.0 μm. A total of 8318 movies (6545 at 0° tilt, 1304 at 20° tilt, 122 at 25° tilt, and 347 at 30° tilt) were collected with the two samples mentioned above and analyzed on CryoSPARC v4.6.0 Live(Punjani et al., 2017) up to 2D classification step.

The cryo-EM data of hAgo2 F294A-guide-target RNA, hAgo2 D669A-guide-target RNA, and hAgo2 WT-guide-target RNA were collected on a TALOS Arctica (Thermo Fisher Scientific) transmission electron microscope with a field emission gun operating at an accelerating voltage of 200 kV. Micrographs were acquired using a Gatan K2 Summit direct electron detector, operated in electron-counting mode, using the automated data collection software Leginon (Suloway et al., 2005) by image shift-based movements from the center of four adjacent holes to target the center of each hole for exposures. Each micrograph was acquired as 23 dose-fractionated movie frames with 4.6 s exposure resulting in a total cumulative electron exposure of 50 e^-^Å^-2^. The datasets were collected with C2 aperture 50 μm, magnification of 45,000x, corresponding to 0.91 Å per pixel on the detector with a nominal defocus range from -1.0 to -1.5 μm. Appion (Lander et al., 2009) image processing was used to run MotionCor2(Zheng et al., 2017) for micrograph frame alignment and dose-weighting in real time during data collection. All subsequent processing steps were performed in CryoSPARC (Fig. S3).

### Image processing and 3D reconstruction

For the processing of hAgo2 R315V/H316A-guide-target RNA data, several steps were performed on CryoSPARC Live, including motion correction, Contrast Transfer Function (CTF) estimation, blob picking, particle extraction, and 2D classification. The 3D reconstruction, refinements, and post-processing were subsequently performed in CryoSPARC. Micrographs with an estimated CTF fit to a resolution of 7 Å or better (7,090 micrographs) were selected for particle picking. A total of 3,286,293 particles were extracted with binning 2×2 (1.48 Å per pixel, 128 pixels box size) and subjected to two rounds of 2D classifications. Based on the similarity to the known Argonaute structure and excluding smaller classes of junk particles, 1,982,073 particles were selected from the 2D class averages.

From the selected particle stack, 100,000 particles were subjected to *ab initio* reconstruction to generate five unique classes. These classes were used for 3D heterogeneous refinement. The best-looking hAgo2 class, chosen from every round, was subjected to the next round of heterogeneous refinement along with a new stack of particles from the collected data in an iterative process. A total of seven rounds of heterogeneous refinements were performed until all the selected particle stacks were processed. The 3D map resembling hAgo2 with a guide-target RNA duplex contained 466,292 particles and was subjected to homogeneous refinement, followed by a local refinement with a mask for the whole complex, followed by another local refinement with a mask around MID, PIWI, and seed region of the complex. A final 2D classification was performed on this particle stack and 9% of the particles were removed as junk. The selected 426,055 particles were subjected to a final round of heterogeneous refinement to remove hAgo2 classes with partial or no guide-target RNA duplex density. The final hAgo2 class with a complete guide-target RNA duplex contained 67% of the particles.

Homogeneous refinement and local refinement were performed on the final particle stack, which was re-extracted with no binning (0.74 Å per pixel, 256 pixels box size) from micrographs with an estimated CTF fit to a resolution of 5 Å or better, and a calculated defocus range of 0.1-2.5 μm. The final 264,433 particles were subjected to homogeneous and local refinement, followed by CTF refinements and final local refinement with a mask of the whole complex. This resulted in a final map of hAgo2 R315V/H316A-guide-target RNA complex with a resolution of 3.16 Å, determined by gold-standard FSC at a cutoff of 0.143 (Fig. S4). Local resolution estimation was performed, and the estimated resolution range was 2.7-5.7 Å. The sphericity and global resolution of the final map were evaluated using the Remote 3DFSC Processing Server (https://3dfsc.salk.edu) (Tan et al., 2017).

For the hAgo2 F294A-guide-target RNA (8323 movies), hAgo2 D669A-guide-target RNA (3106 movies), and hAgo2-guide-target RNA (1847 movies) data processing, motion-corrected micrographs were subjected to patch CTF estimation, followed by blob picking and 2D classification. For the F294A mutant sample, six *ab initio* classes were generated. For WT and D669A samples, initial *ab initio* maps were generated from previously screened cryo-EM maps of the hAgo2 WT-guide-16nt target RNA complex. The *ab initio* maps were used to run heterogeneous refinements iteratively for selected particle stacks. The best class from the last round of heterogeneous refinement was subjected to non-uniform refinement, followed by local refinement.

For hAgo2 F294A-guide-target RNA, two final conformations were reconstructed. Conformation 1, with 88,631 particles and a resolution of 4.75 Å, was reconstructed to a map of hAgo2 with a complete guide-target RNA duplex. Conformation 2, with 58,244 particles and a resolution of 6.47 Å, was reconstructed to a map of hAgo2 with guide-only conformation (Fig. S10).

### Model building and refinement

For hAgo2 R315V/H316A-guide-target RNA model building, the initial model of hAgo2 (no RNA) was obtained from the crystal structure of hAgo2 bound to a seed and supplementary paired target (PDB ID: 6N4O). The guide-target RNA duplex was built and modeled using Coot (Emsley et al., 2010). The domains of hAgo2 and the guide-target RNA duplex were first fit into the final cryo-EM map as a rigid body in ChimeraX (Goddard et al., 2018; Meng et al., 2023), followed by a rigid body refinement in Phenix (Liebschner et al., 2019) to generate an initial model. The real-space refinement of the initial model was performed by optimizing global minimization, atomic displacement parameters, and local grid search using Phenix. Iterative rounds of manual building and fixing geometric, rotamer, and Ramachandran outliers were performed in Coot and ISOLDE (Croll, 2018) in ChimeraX. The final model was validated by comprehensive validation (cryo-EM) and EMRinger(Barad et al., 2015) suite in Phenix. The PAZ domain (residue 228-347) along with a few other residues left unmodeled due to a lack of interpretable density. RNA coordinates where analyzed using Web 3DNA 2.0 (Li et al., 2019).

