## Supplementary Figures and Table for "Structural basis for gene silencing by siRNAs in humans"

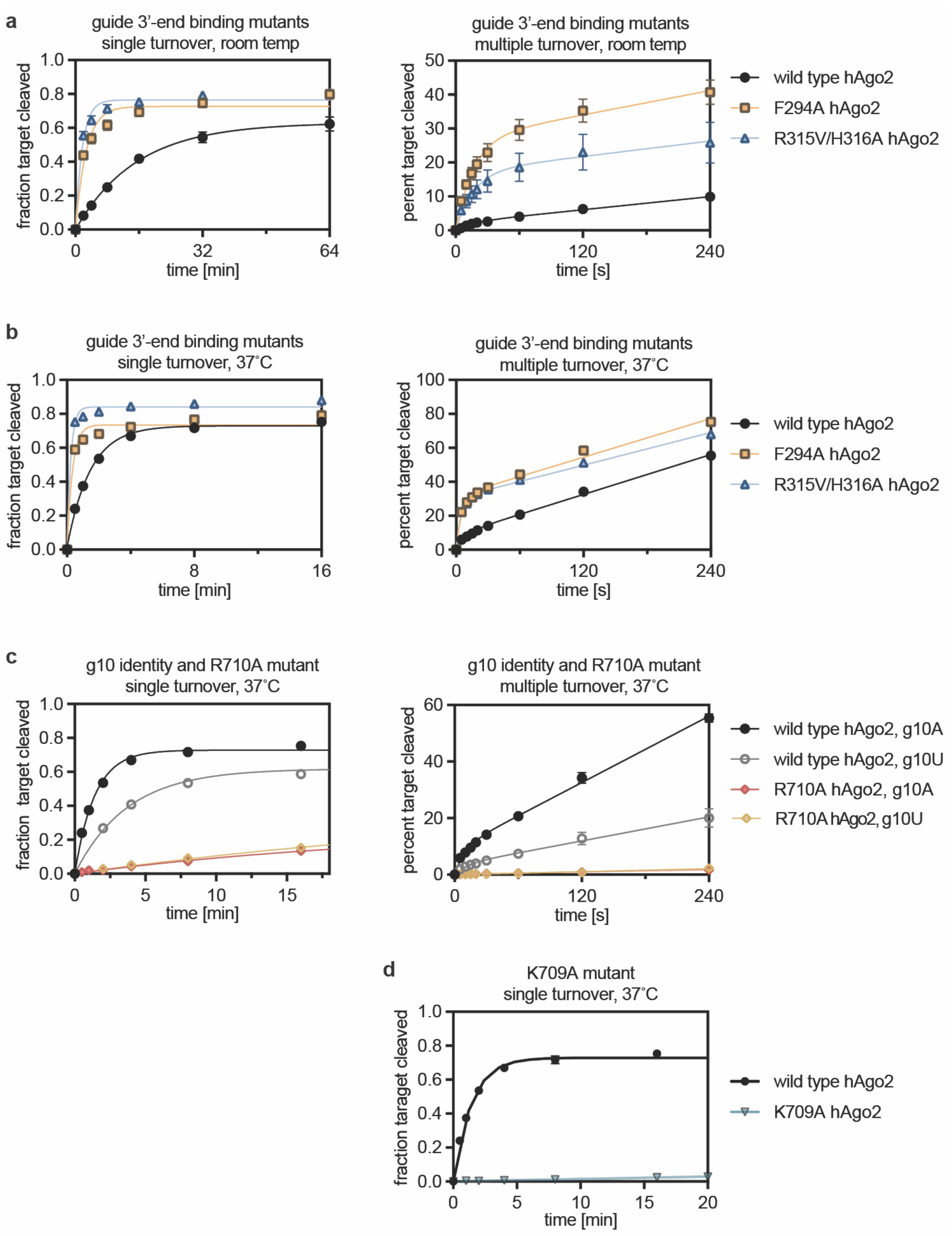


**Supplementary Figure 1: Kinetic target cleavage data in this study. a.** Target cleavage under single turnover (left) and multiple turnover (right) conditions by guide 3'-end binding mutants at room temperature, matching the conditions used to prepare cryo-EM grids. **b.** Target cleavage under single (left) and multiple (right) turnover conditions by guide 3'-end binding mutants at 37˚C. **c.** Target cleavage under single (left) and multiple (right) turnover conditions by wild type and R710A hAgo2 using guides with g10A or g10U nucleotides at 37˚C. Target cleavage under single turnover conditions by the K709A hAgo2 mutant at 37˚C. All points represent the mean value from 3 experiments. Error bars, when visible, indicate SEM.

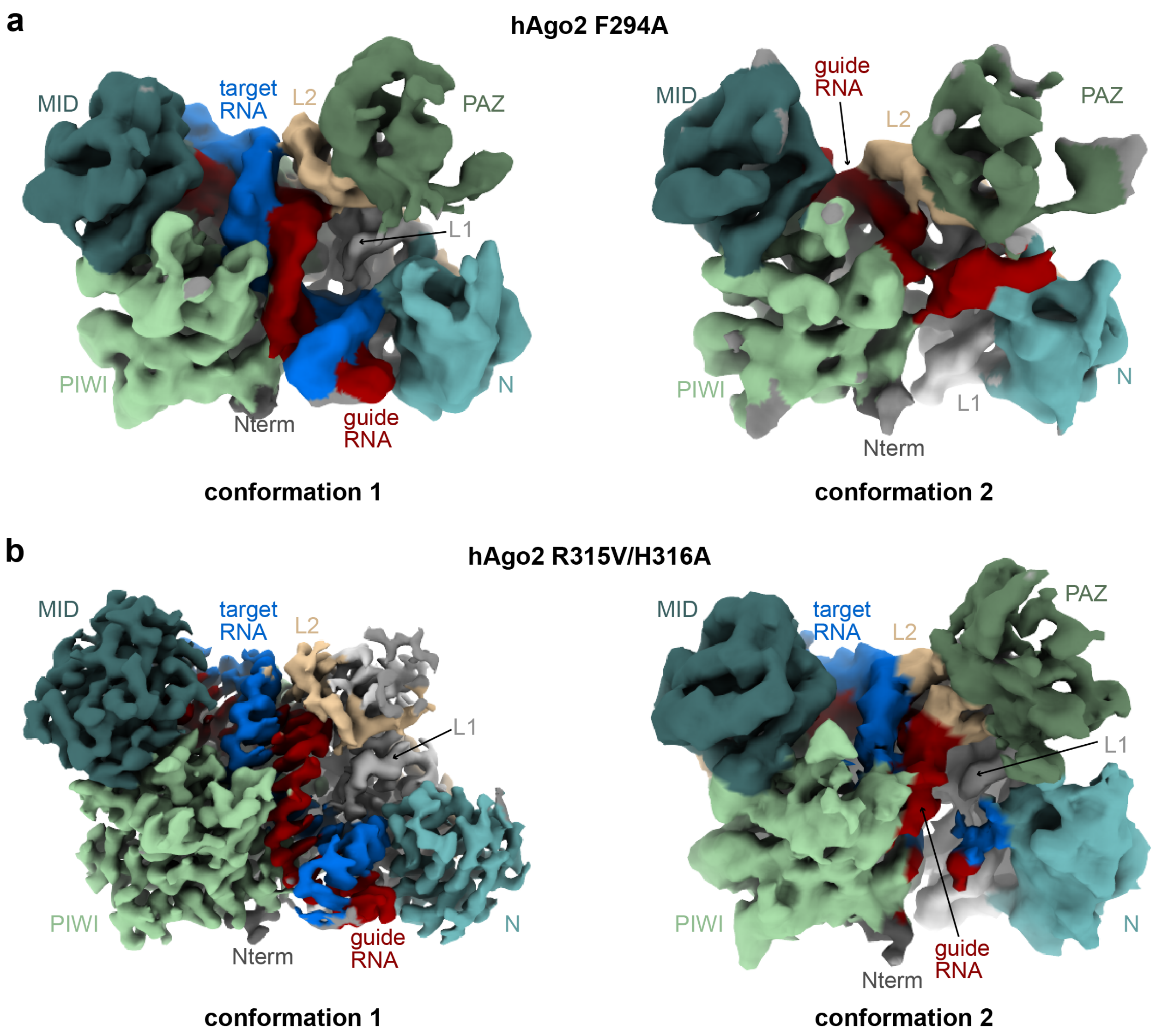


**Supplementary Figure 2: Reconstructions of guide 3'-end binding pocket mutant hAgo2 proteins engaged with guide and fully complementary target RNA.** Both the F294A and R315V/H316A Ago2 mutants produced reconstructions with density for an intact guide-target duplex in the central cleft (left). Alternate conformations were observed as well (right). Data for the R315V/H316A Ago2 mutant were higher quality and thus used for high-resolution reconstruction and analysis.

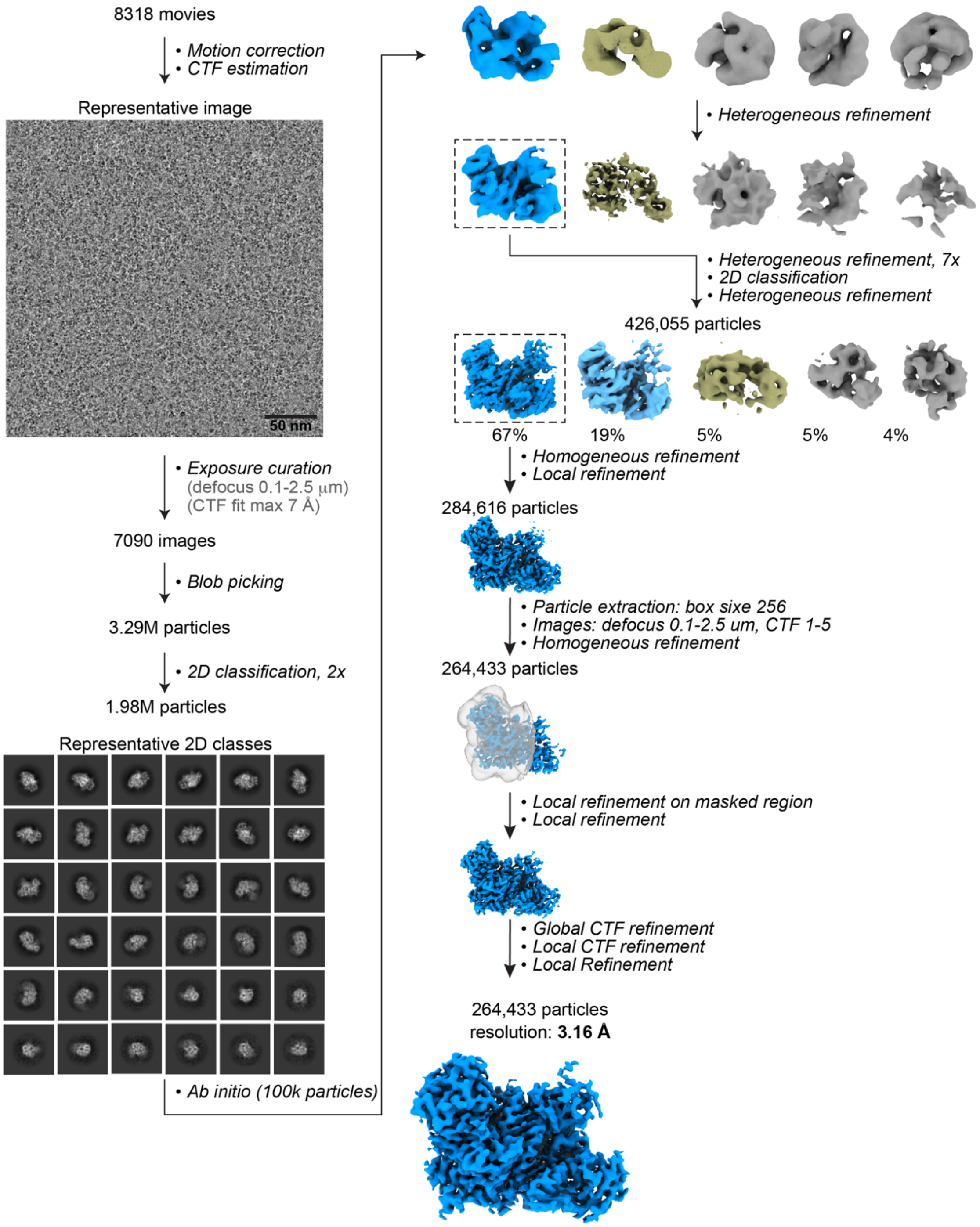


**Supplementary Figure 3: Cryo-EM Imaging and processing of the hAgo2 R315V/H316A-guide-target RNA complex.** Representative data processing workflow. Motion correction, CTF corrections, blob picking, extraction, and 2D classification were performed in CryoSPARC live. Selected particles with high-resolution features were exported for further 3D processing. First, 100,000 particles were used to create initial Ab initio 3D maps. These maps were used as input 3D volumes for heterogeneous refinement of all the selected particles. A total of 8 rounds of heterogeneous refinements were performed. The class of particles with the highest resolution and complex-like features from each round were chosen for further refinement until the other noisy low-resolution classes were less than 10%. The class with the highest resolution with complex-like features was chosen from the final heterogeneous refinement for further refinements and post-processing. The final 3D volume of hAgo2-miR-122 in complex with a fully complementary target was reconstructed from 264,433 particles, achieving a resolution of 3.16 Å.

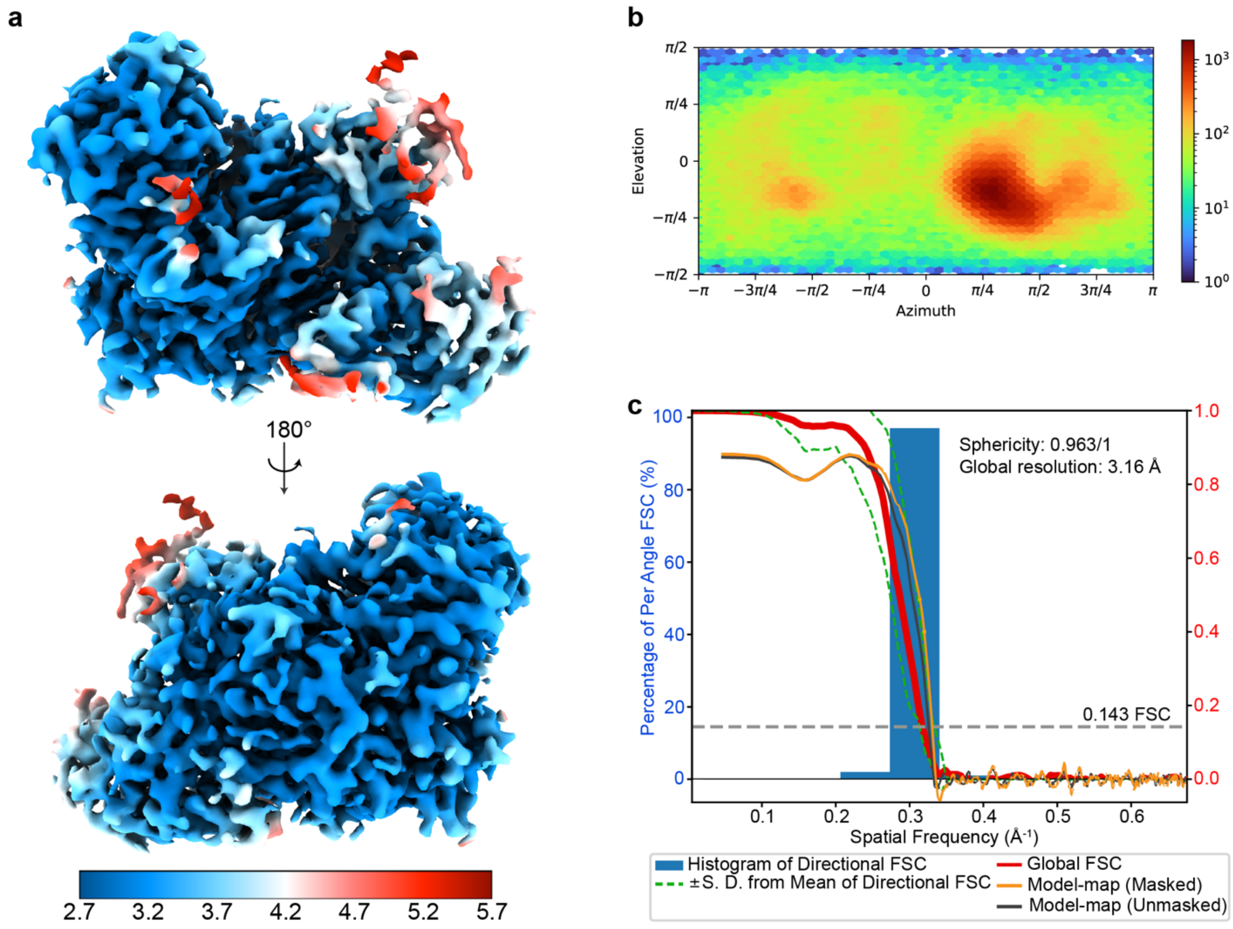


**Supplementary Figure 4: Quality of the cryo-EM reconstruction of hAgo2 R315V/H316A-guide-target RNA complex. a.** Local resolution representation of the final map. The map is colored according to local resolution estimation. **b.** Angular distribution plot of the final map of the complex. **c.** A 3DFSC histogram with map-to-model FSC is overlaid.

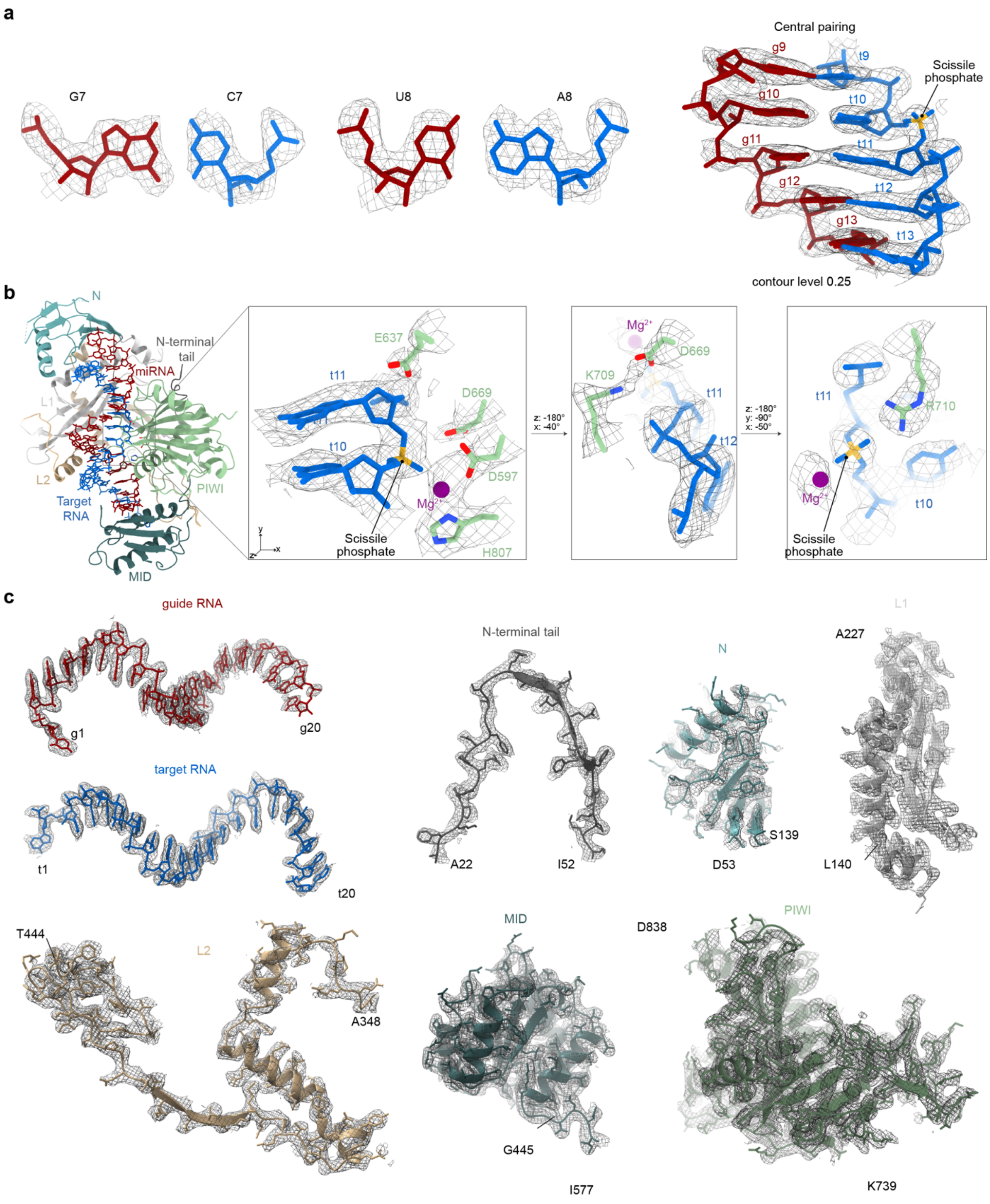


**Supplementary Figure 5**: **Map quality. a.** RNA density quality was often high enough to unambiguously visualize nucleobase identity. Map quality in the central region (right), which contains the scissile phosphate and an extended major groove. **b.** Map quality of the previously established catalytic tetrad (D597, E637, D669, and H807) and the residues identified in this study: K709, proposed to facilitate protonation of the leaving group, and R710, proposed to stabilize charge on the transition states. **c.** Density for individual domains of hAgo2 and RNAs fit into the cryo-EM map. The map is shown as a mesh; protein models are shown in cartoon representation with side chains as sticks; RNAs are shown in stick representation.

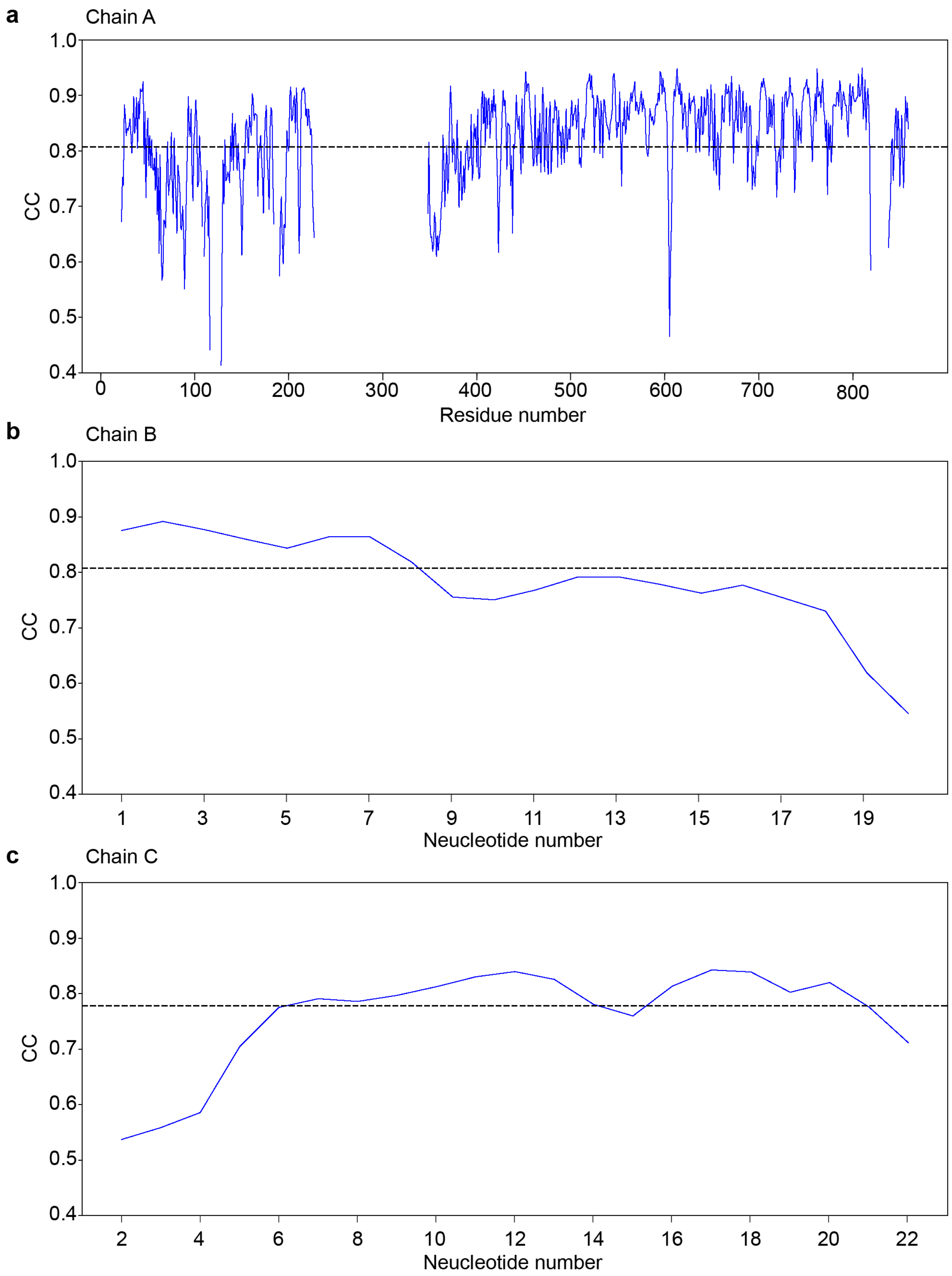


**Supplementary Figure 6: Correlation Coefficient plots for the model of the hAgo2 R315V/H316A-guide-target RNA complex.** Model refinement was performed in Phenix. Per-residue Correlation Coefficient (CC) plots for **a.** the protein chain, **b.** miR-122 21 mer guide RNA, and **c.** FT21 target RNA are shown. Solid blue lines represent per-residue CC and dashed gray lines represent average CC for the chain.

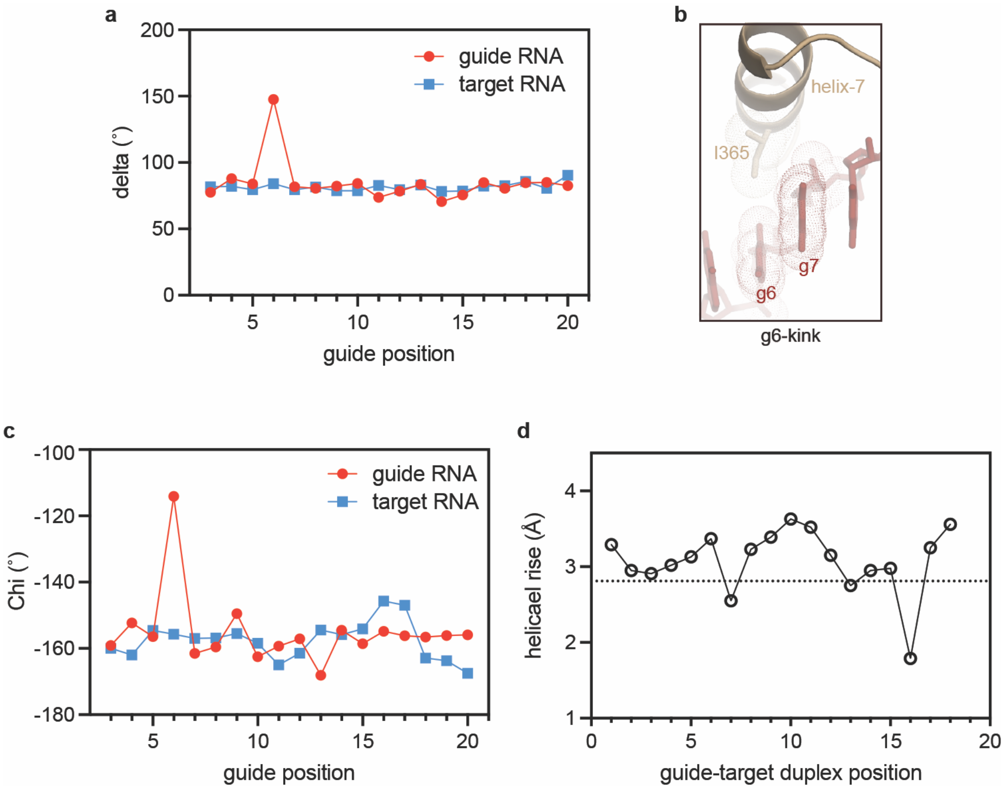


**Supplementary Figure 7: Helical distortion data. a.** Plotting the delta (δ) torsion angle, which describes sugar pucker, for all nucleotides in the guide-target duplex shows only g6 adopts the C2' endo conformation. **b.** Hydrophobic interactions between Ile-365 and the sugar face at positions 6-7 connect the guide-target duplex to helix-7. **c.** Plotting the Chi (χ) torsion angle, which describes the relationship of the ribose and nucleobase, for guide and target nucleotides shows local distortions in the guide strand at the g6-kink and the target strand at positions 16-17, associated with compression of the major groove. **d.** Plotting helical rise between stacked bases reveals local duplex compressions at the g6-kink and position 16.

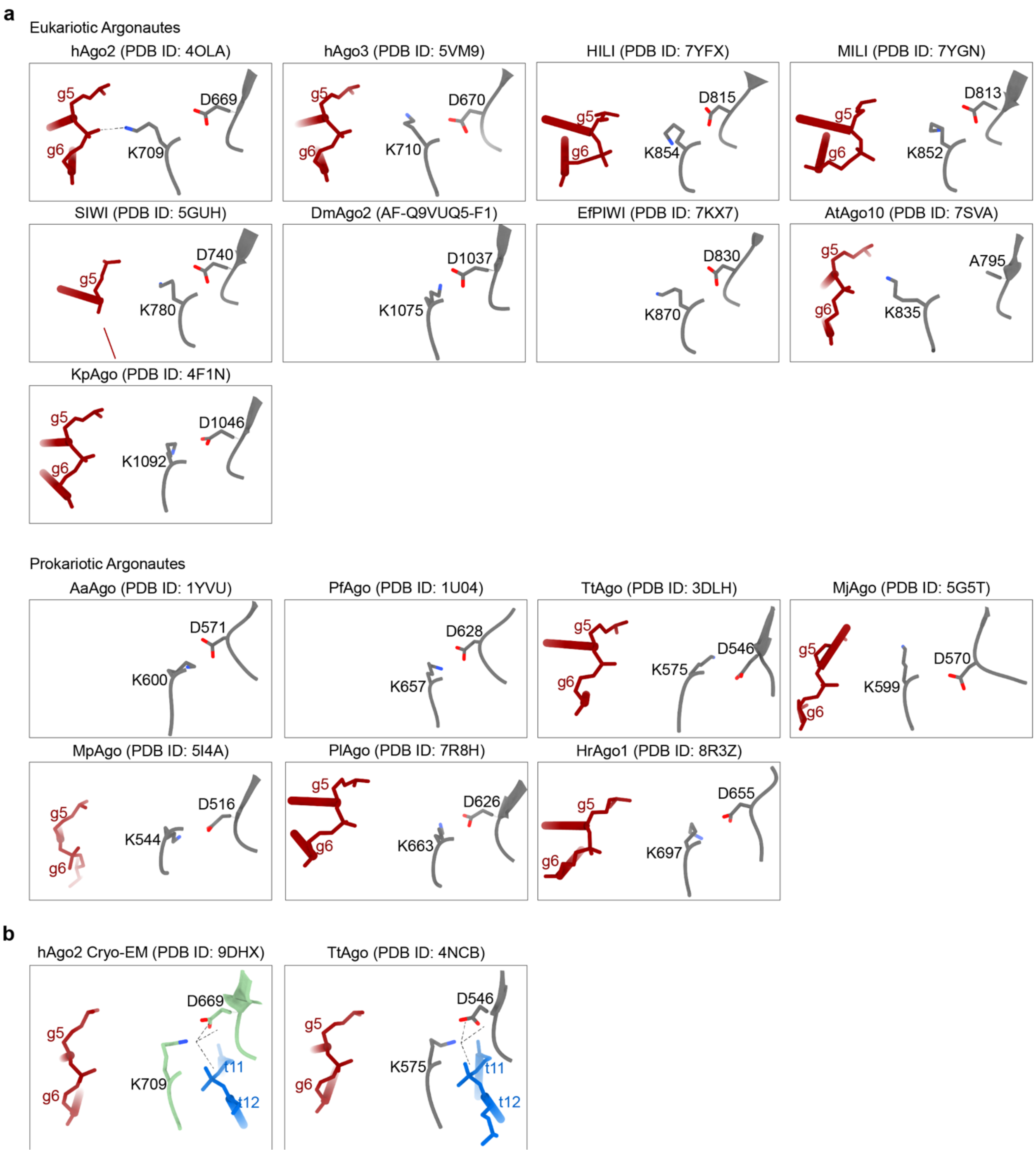


**Supplementary Figure 8: The Asp-Lys pair is a deeply conserved feature of the Argonaute active site. a.** Asp-Lys pairs observed in apo and guide RNA bound Argonaute protein family structures. hAgo2 appears to be special in that the Lys residue in most Ago proteins interrogated does not interact with the guide RNA. **b.** the structures of Argonautes with guide-target RNA duplex. Comparing TtAgo structures with and without target RNA suggests the formation of the Asp-Lys pair is target RNA dependent.

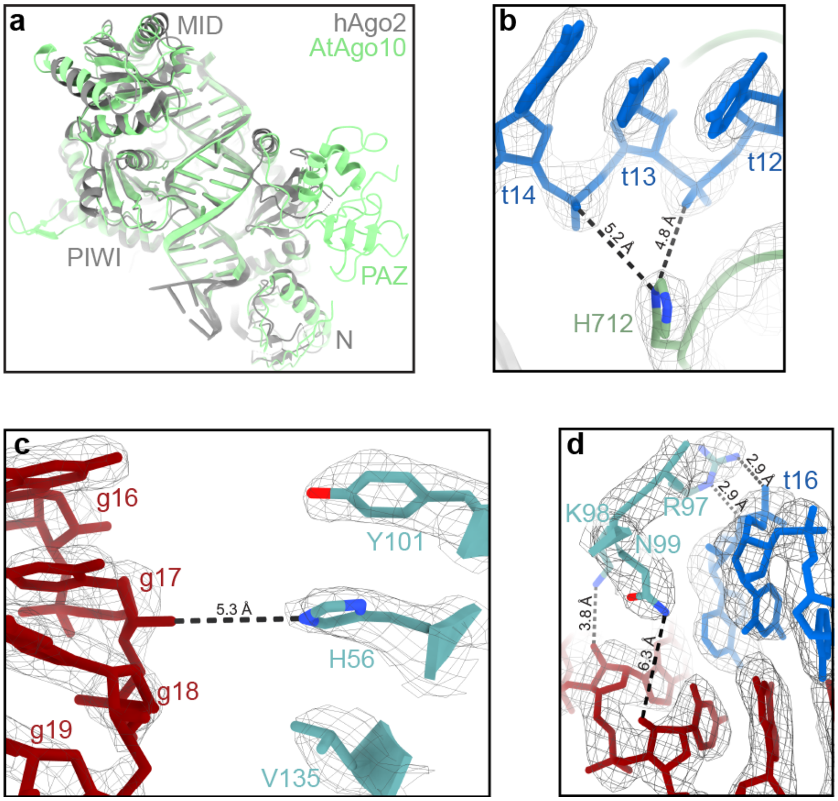


**Supplementary Figure 9. Differences between reported hAgo2 catalytic structures.** Differences with functional implications between the hAgo2-guide-target structure recently reported by Mohamed, *et al.* and the structure presented here are described below:

**a.** Mohamed, *et al.* reported that the N domain of hAgo2 shifts ~7 Å compared to the N domain position in the AtAgo10-guide-target structure, which had a guide target duplex ending at position 16, and concluded that this shift is critical for allowing the full RNA duplex access to the central channel. Although our hAgo2 structure contains a full RNA duplex (to position 21), we do not observe this shift. Specifically, when the MID-PIWI lobes of our hAgo2 (grey) and AtAgo10 (green) are aligned, the respective N domains fall within 2.3 Å of each other (based on both center of mass displacement and the average distance between all equivalent Ca atoms). This difference might be attributed to a shift induced by the longer siRNA-target duplex used in the hAgo2 structure, as proposed by Mohamed, *et al.* However, alternative explanations, such as differences in sequence and structure of plant and human Ago proteins, or inaccurate modeling of the N domain in the lower-resolution AtAgo10 map cannot be excluded. More importantly, we do not observe a clash with the hAgo2 RNA duplex modeled onto AtAgo10. We conclude that an exaggerated N domain shift (beyond the shift observed for AtAgo10 previously) is not a strict requirement for full RNA access to the hAgo2 central cleft and that, in some cases, compression of the guide-target duplex instead enables RNA docking.

Panels b-d show how indicted regions of our structure fit in the experimental map (wire mesh) with distances between proposed interacting atoms indicated:

**b.** Mohamed *et al.* proposed that H712 of the ‘bridge loop’ in the PIWI domain interacts with backbone phosphates of target nucleotides t12 and t13 to help position centrally paired RNA in the active site for slicing. In our hAgo2 structure, H712 is too far from the t12-t13 phosphates to establish direct interactions.

**c.** Mohamed *et al.* proposed that H56 of the N domain contacts RNA backbone phosphates to favor slicing by stabilizing the slicing-competent conformation. H56 is recessed at the hAgo2 surface, sandwiched between hydrophobic residues Y101 and V135, and is not positioned to contact the RNA in our structure.

**d.** Mohamed *et al.* proposed that R97, K98, and N99 stabilize the slicing-competent conformation through interactions with the RNA backbone phosphates. Of these three residues, only R97 contacts a backbone phosphate in our model.

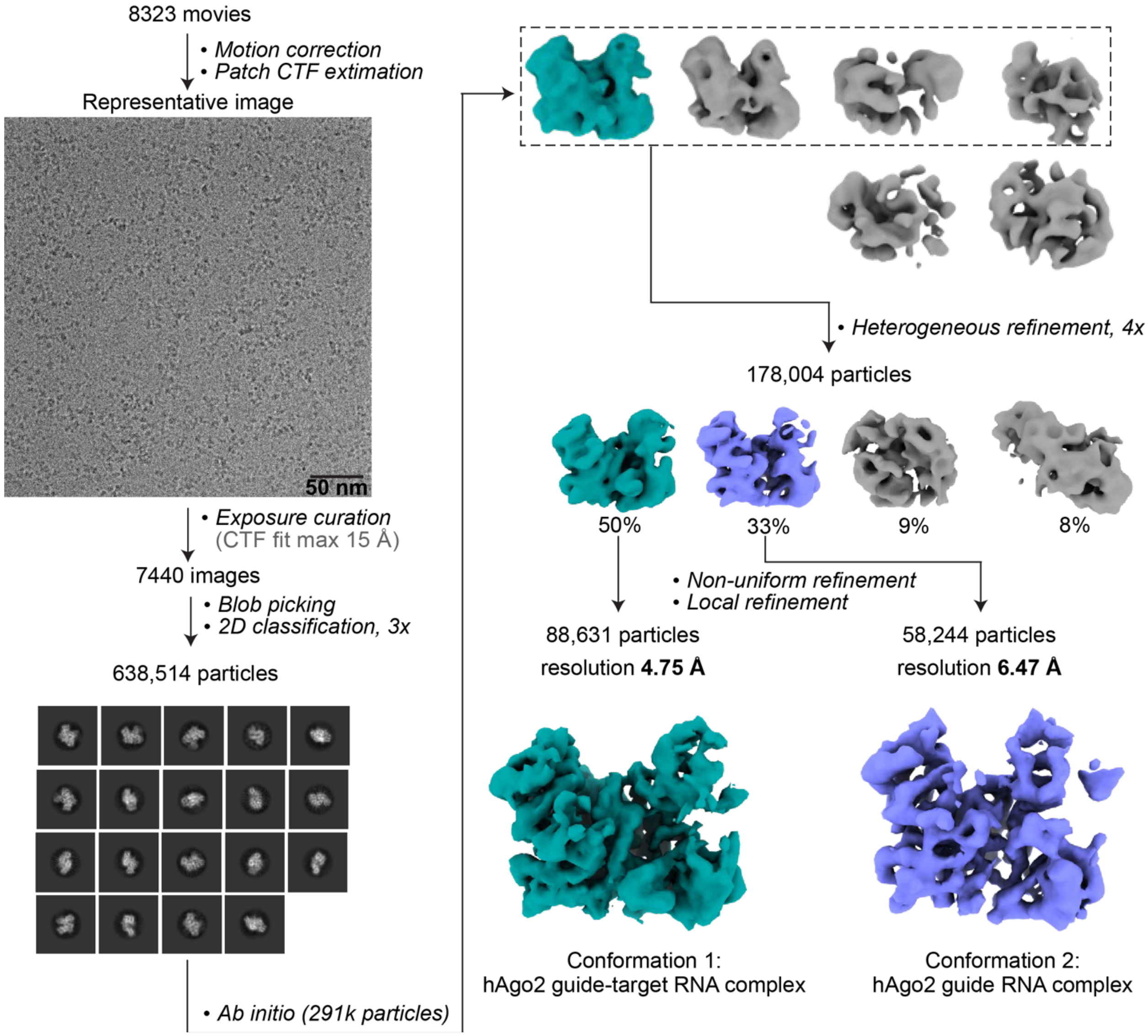


**Supplementary Figure 10: Cryo-EM Imaging and processing of the hAgo2 F294A-guide-target RNA complex.** Representative data processing workflow. Motion correction, CTF corrections, blob picking, extraction, and 2D classification were performed in CryoSPARC live. Selected particles with high-resolution features were exported for further 3D processing. First, 291,000 particles were used to create initial Ab initio 3D maps. The ab initio maps were used to run heterogeneous refinements iteratively for selected particle stacks. The best class from the last round of heterogeneous refinement was subjected to non-uniform refinement, followed by local refinement. For hAgo2 F294A-guide-target RNA, two final conformations were reconstructed. Conformation 1, with 88,631 particles and a resolution of 4.75 Å, was reconstructed to a map of hAgo2 with a complete guide-target RNA duplex. Conformation 2, with 58,244 particles and a resolution of 6.47 Å, was reconstructed to a map of hAgo2 with guide-only conformation

**Supplementary Table 1: Cryo-EM data collection, refinement, and validation statistics of hAgo2 R315V/H316A-guide-target RNA complex**

| | |
|---|---|
| (EMDB-46888)<br>(PDB 9DHX) | |
| **Data collection** | |
| Microscope | Talos Arctica |
| Camera | FEI Falcon |
| Camera mode | counting |
| Magnification | 190,000 |
| Voltage (kV) | 200 |
| Total electron exposure ($e^-/Å^2$) | 40 |
| Exposure rate ($e^-$/pixel/s) | 6.83 |
| Exposure time (s) | 3.2 |
| Defocus range (μm) | -1.0 to -3.0 |
| Pixel size (Å) | 0.74 |
| Symmetry imposed | C1 |
| Data acquisition software | EPU |
| Micrographs collected (no.) | 8318 |
| **Data analysis** | |
| Total extracted particles (no.) | 3,286,293 |
| Particles used for 3D analyses (no.) | 1,982,055 |
| Final refined particles (no.) | 264,433 |
| Symmetry | C1 |
| Global resolution (Å) | |
| FSC 0.5 (unmasked / masked) | 3.9 / 3.4 |
| FSC 0.143 (unmasked / masked) | 3.6 / 3.2 |
| Local resolution range (Å) | 2.7 - 5.7 |
| 3DFSC Sphericity (%) | 96.3 |
| Map sharpening *B*-factor ($Å^2$) | 118.5 |
| **Model composition** | |
| Chain | 4 |
| Non-hydrogen atoms | 6312 |
| Protein residues | 683 |
| Nucleotides | 41 |
| Ligands ($Mg^{2+}$ ion) | Mg: 1 |
| **Model Refinement** | |
| Refinement Package | Phenix/ISOLDE |
| MapCC (volume/mask) | 0.83/0.86 |
| *B*-factors ($Å^2$) | |
| Protein residues | 58.90 |
| Nucleotides | 61.88 |
| Ligand ($Mg^{2+}$ ion) | 30.00 |
| R.m.s. deviations | |
| Bond lengths (Å) | 0.013 |
| Bond angles (°) | 1.837 |
| **Validation** | |
| Map-to-model FSC 0.5 (unmasked/masked) | 3.3 / 3.2 |
| Map-to-model FSC 0.143 (unmasked/masked) | 3.1 / 3.0 |
| Ramachandran plot | |
| Favored (%) | 96.42 |
| Allowed (%) | 3.58 |
| Outliers (%) | 0.00 |
| MolProbity score | 0.99 |
| Clashscore | 0.82 |
| Poor rotamers (%) | 0.00 |
| CaBLAM outliers (%) | 2.43 |
| EMRinger score | 4.99 |
